# Engineered histones reshape chromatin in human cells

**DOI:** 10.1101/2025.09.10.674980

**Authors:** Siddhartha G. Jena, Surya Nagaraja, Andrew S. Earl, Amalia R. Driller-Colangelo, Michael A. Quezada, Ena Oreskovic, Max A. Horlbeck, Ruochi Zhang, Wilson Gomarga, Jason D. Buenrostro

## Abstract

Histone proteins and their variants have been found to play crucial and specialized roles in chromatin organization and the regulation of downstream gene expression; however, the relationship between histone sequence and its effect on chromatin organization remains poorly understood, limiting our functional understanding of sequence variation between distinct subtypes and across evolution and frustrating efforts to rationally design synthetic histones that can be used to engineer specified cell states. Here, we make the first advance towards engineered histone-driven chromatin organization. By expressing libraries of sequence variants of core histones in human cells, we identify variants that dominantly modulate chromatin structure. We further interrogate variants using a combination of imaging, proteomics, and genomics to reveal both *cis* and *trans-*acting mechanisms of effect. Functional screening with transcription factor libraries identifies transcriptional programs that are facilitated by engineered histone expression. Double mutation screens combined with protein language models allow us to learn sequence-to-function patterns and design synthetic histone proteins optimized to drive specific chromatin states. This work establishes a foundation for the high-throughput evaluation and engineering of chromatin-associated proteins and positions histones as tunable nodes for understanding and modulating mesoscale chromatin organization.

## Introduction

Cells coordinate proper gene expression and responses to their environment by modulating chromatin structure, balancing DNA accessibility with protection from damage. To manage this tradeoff, eukaryotes use histone proteins to shape the physical and regulatory landscape of the genome by packaging DNA into nucleosomes [1–2]. At individual genes, nucleosomes occlude the binding of transcription factors (TFs), while at the genomic level, chromatin state modulates cell plasticity and safeguards DNA. Chromatin’s structural diversity arises from post-translational modifications (PTMs) [3–5] as well as the incorporation of histone variants: specialized paralogs that can profoundly alter genome organization and gene expression [6].

Across nature, diverse organisms have evolved specialized histone variants to ensure diverse physical properties of chromatin enabling control across tissues [7–8], viral invasion [9], and host defense [10] (**Figure 1A**). These variants include genetic mutations that mimic the effect of histone PTMs or alter DNA-protein interactions to alter chromatin structure. Indeed, prior studies have used variants and gene mutations mimicking histone PTMs to achieve significant gains in the efficiency of cell reprogramming [7, 11] or combating senescence [12]. These findings highlight the regulatory potential encoded in histone sequence and suggest that histones could be further engineered to diversify the regulatory function of the genome; for example, opening heterochromatin regions to enable cellular plasticity or increasing the responsivity of the genome to transcription factors for cellular reprogramming. However, this suite of potential applications are limited by the incomplete understanding of the sequence-to-function relationship of histones.

**Figure 1:**
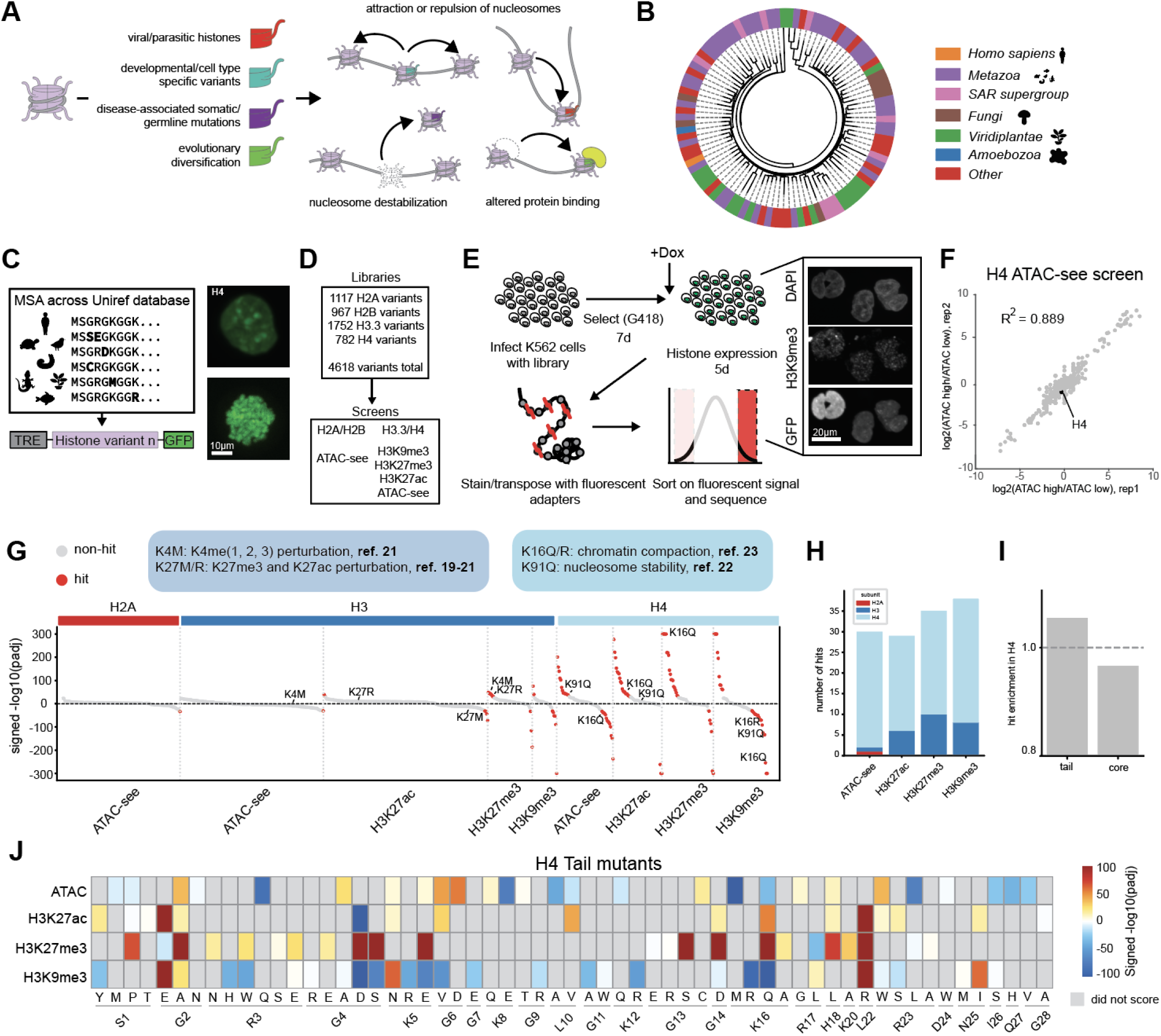
A high-throughput screen of histone mutations reveals modulators of chromatin structure. A. histone variants are deployed throughout nature to modulate chromatin in a range of contexts, and utilize several mechanisms to accomplish their tasks. B. Diversity of the H4 protein family across phyla. Outer leaves are protein sequence clusters of all 102-aa length (same length as *Homo sapiens* H4C1) H4 proteins on the UniRef database. C. Building a histone mutant library in an inducible and GFP-tagged expression vector (left) and confirming proper expression (right, top image) and integration into chromatin (right, bottom image depicting GFP in mitotic chromatin). D. Variant libraries constructed and screens performed. E. Workflow for histFlow screens. Inset image is of example stained cells imaged in 3 colors corresponding to DNA staining, GFP-tagged histones and H3K9me3 staining. F. Correlation between replicates of the H4 ATAC-see screen. G. Significant hits (defined by padj < 0.05) from all screens, separated by histone subunit (top label) and then screen (bottom label). Red dots denote hits with-log10(padj) > 20. Mutants that arise in previous literature are labeled. H. Enumeration of hits in each type of screen, by histone subunit. I. enrichment of hits in the H4 tail vs. globular domain, normalized to the length of the domain. J. H4 tail mutant hits, colored by signed log(padj). Gray boxes denote that the mutant was not a significant result of the given screen.

To address this challenge, we have generated a lentiviral library of 4,618 histone variants spanning the core nucleosomal subunits H2A, H2B, H3, and H4. Using this library, we were able to perform large-scale screens of mutant histones on global chromatin accessibility and histone modification state (histFlow). Interestingly, we find that the H4 histone tail is enriched for mutations that disrupt heterochromatin and alter global chromatin structure. We highlight the H4G4D mutant, which disrupts heterochromatin in *cis* and *trans,* alters spatial compartmentalization within the nucleus, and reconfigures the functional TF binding landscape in T cells. Finally, by incorporating multiple-mutation histFlow data, we design and train a model architecture (histForge) that predicts the effect of histone sequence variants on heterochromatin abrogation and accessibility change. With this approach, we design synthetic histone variants that de-repress heterochromatin. Our work demonstrates that engineered histone variants can reprogram genome-wide chromatin state and provides a framework for the design of chromatin state in living cells.

### histFlow screens reveal modulators of global chromatin organization from a library of evolutionary sequence variants

Our work is inspired by previous work that seeks to describe the functional consequences of histone mutations [13–14]. Prior studies, in particular those motivated by mutations associated with human disease and variants associated with developmental programs and pathogenesis, have described histone variants with mechanisms that block or structurally mimic the deposition of particular PTMs to promote altered chromatin states [13], while others alter the dynamics or kinetics of nucleosome turnover [14], causing increases or decreases in accessibility of the genome to regulatory proteins (**Figure 1A**). Motivated by these foundational experiments, we sought to catalogue the genetic diversity of histone sequences to ultimately define a gain-of-function library of genetic variants. Accordingly, we turned to evolution to build a database of select variants that had a high probability of being compatible with life. We first sought to characterize the diversity of the histone family across evolutionary space by constructing Hidden Markov Model (HMM)-based alignments [15] between *Homo sapiens* histone proteins and a total of 128,987 histone homologs from the UniRef database [16]. Although histones are relatively more conserved than many other protein families, we still found extensive diversity, revealing 3,086 unique histone sequences that were the same length as the corresponding *Homo sapiens* sequences (**Figure S1A**). Clustering only these sequences (for H2A, H2B. H3.3, and H4) and labeling the phyla composition of the 50 largest clusters revealed sequence trends that are conserved between subsets of metazoans, fungi, plants, and other phyla (**Figure 1B, S1B**).

We used the human-length subset of each histone protein to call individual amino acid variants, creating a library spanning single mutations which were putatively compatible with life. This analysis revealed sites that seemed to tolerate more or less variability; for example, tail regions, which are characteristically disordered regions, had higher sequence diversity than globular regions, which likely have stronger structural constraints in order to wrap DNA. As a result, representation across generated libraries generally reflected the relative conservation of residues at the given position (**Figure S1C**). Variants were cloned into a lentiviral backbone (see **Methods**) and tagged with GFP to monitor nuclear localization for downstream imaging applications; this additionally allowed us to confirm that our construct was properly integrating into both interphase and mitotic chromatin (**Figure 1C**). Histone expression was also placed under doxycycline-inducible control, to precisely control expression duration (**Figure S1D**). Based on this construction, we reasoned that in contrast to most gene mutagenesis screens, which often seek to demarcate loss of function variants, our library would likely be enriched for gain of function properties and depleted in variants yielding dysfunctional, potentially cytotoxic proteins.

We screened our histone libraries on the total levels of three distinct histone modifications: H3K27ac, H3K27me3, and H3K9me3, which describe distinct euchromatin and heterochromatin states. We also used chromatin accessibility, measured by transposase-based profiling and imaging (ATAC-see, **Figure S2A, S2B**), as an additional readout [17–18]. Together, these approaches, which we named histFlow, enabled a systematic, high-throughput exploration of how histone sequence variants remodel chromatin at both structural and regulatory levels. ATAC-see histFlow screens were performed for all histone libraries, and epitope histFlow screens were performed for H3.3 and H4 (**Figure 1D**). For histFlow screening, histone libraries were expressed for 5 days in K562 cells, which are commonly used for both screening and epigenomics studies, then subjected to transposition or epitope staining, prior to flow cytometry and downstream sequencing (**Figure 1E**). The top and bottom 10% of cells were collected based on total accessibility and sequenced to identify mutants (see **Methods**). Screens demonstrated concordance between replicates, with most R^2^ values between 0.88 and 0.92 (**Figure 1F, Figure S2C**). Importantly, concurrent viability screens conducted on doxycycline treated and untreated cells suggested that histone overexpression did not significantly alter cell fitness, suggesting that the variants chosen were in fact compatible with cellular function on the timescale over which the screen was conducted (**Figure S2D**).

Our screens revealed hotspots of functional chromatin alteration. Plotting significantly enriched or depleted mutants (signed FDR < 0.05) revealed some slight effects in H2A mutants, and a number of effects in both H3 and H4 (**Figure 1G**). Some of these hits have been previously implicated in changing chromatin state (**Figure 1G, labeled**): for instance, H3.3K27M and K27R, which disrupt H3K27 methylation and acetylation and appeared as hits in those respective screens [19–21]. H3K4M also appeared as a hit in our H3K27me3 screen, consistent with its reported effects on methylation/acetylation at that site in addition to its proposed effect on H3K4me [21]. Hits also included mutants known to be drivers of disease, such as Tessadori-Bicknell-van Haaften (TBV) mutations in H4 (R40C, R45C, and K91Q, all of which caused increases in accessibility in accordance with their proposed mechanism of action [22]). Finally, mutations of H4K16 (K16Q and K16R), which were also recovered in our screens, mimic acetylation states of K16, a residue that has been investigated in prior work and found to have significant effects on chromatin compaction *in vitro* by inhibiting internucleosomal interactions [23]. These mutants can, when overexpressed, lead to organismal effects; for instance, extending lifespan in yeast [12]. Most of these established mutations occur at lysines and are thought to operate through mimicking or disrupting PTMs. However, our screens also recovered some unexpected patterns of behavior: most dramatically, a significant number of sequence variants inducing chromatin changes within H4 (**Figure 1H**). In particular, the tail region of H4, despite its small size and little direct interaction with wrapped DNA, was slightly enriched for hit mutants (**Figure 1I, J**). Most interestingly, many of these mutants occurred at non-lysines: for example, the H4G4D mutant, which occurred at a glycine that is not thought to be modified. In particular, the ability of some H4 tail mutants to reduce H3K9me3 levels was intriguing, as this mark denotes constitutive heterochromatin and is considered to be a major barrier to TF binding and cell state changes [24]. We validated a subset of tail mutants in stably expressing K562 lines (**Figure S2E-G**), consisting of both established (e.g. K16Q, K16R) and new (e.g. G4D) mutants that affected chromatin structure. We decided to investigate these mutants further, as there has been recent appreciation of the role of histone tails in and the H4 tail in particular for their roles in modulating TF binding and chromatin compaction [25–26].

### Spatial characterization of H4 tail mutants reveals *cis* and *trans* effects on heterochromatin

We next sought to understand how H4 tail mutants nominated by histFlow interact with and affect heterochromatin. One possibility was that histones worked in *cis* to destabilize chromatin structure at the integrated nucleosome site. Alternatively, nucleosomes may work in *trans. Trans* mechanisms are diverse [19–20,27], including mechanisms wherein nucleosomes sequester chromatin readers and writers from their endogenous sites. Indeed, both of these mechanisms have been reported in prior literature [19–20,27] (Fig. 1A). We reasoned that although both *cis* and *trans* effects could reduce H3K9me3 levels, a *cis* effect would lead to changes in H3K9me3 near the site of histone incorporation, while a *trans* effect would likely lead to non-localized changes. To test this, we used imaging of stable mutant cell lines for H3K9me3 and, using an automated high-content imaging platform (see **Methods**), imaged both heterochromatin and histone-GFP distribution. We noted a range of GFP localization behavior: while wildtype H4 demonstrated some colocalization between histone-GFP and H3K9me3 (**Figure 2A**), some mutants, such as G4D (**Figure 2B**) displayed strong exclusion of GFP from heterochromatin foci. Even the H3K9M mutant, which is known to decrease H3K9me3 levels [27–28], did so, but with no change in physical distribution of GFP, consistent with its proposed *trans-*acting mechanism (**Figure 2C**). Performing this analysis across thousands of cells (1,500-2,000 cells per mutant) confirmed this observation, as total H3K9me3 levels decreased for all examined mutants (**Figure 2D**), while a select few (G4D, K16R, G14D, R3H, and K16Q) demonstrated significant exclusion from heterochromatin **(Figure 2E**). Mitotic chromatin profiling confirmed that this effect was not indirectly caused by lack of integration into chromatin (**Figure S3A**). Additionally, GFP CUT&Tag performed on a selection of mutants showed little change in the distribution of GFP-tagged histones in H4-GFP and tail mutants tested, suggesting that they were in fact deposited in similar patterns (R^2^ > 0.92 for all comparisons, **Figure S3B**).

**Figure 2:**
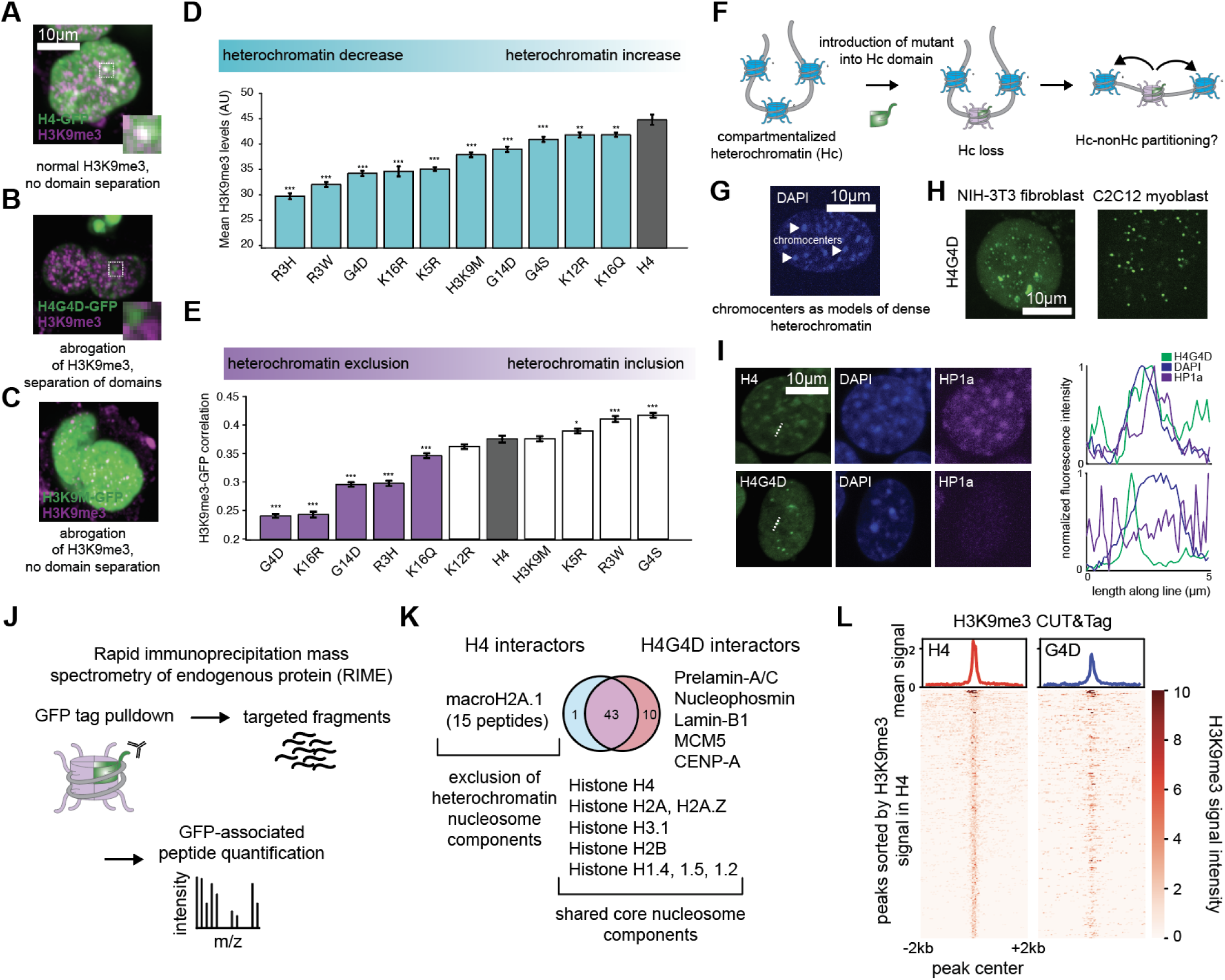
**H4 tails destabilize heterochromatin and alter spatial compartmentalization in *cis* and *trans.*** A-C. Differences in colocalization of GFP-tagged histones with H3K9me3, shown for H4, G4D, and K9M. D. Total H3K9me3 staining in hits from the H3K9me3 screen, showing consistent decrease in H3K9me3 levels. E. Correlation between per-pixel GFP and H3K9me3 signal in hit cell lines; certain hits from the H3K9me3 screen display significant exclusion from H3K9me3 signal. (*n* = 2502-5386 cells per mutant for D and E, with ** corresponding to *p* < 0.05 and *** corresponding to *p* < 0.005. Error bars represent SEM) F. Schematic outlining proposed mechanism of mutant histone (green) incorporating into nucleosome and resulting in partitioning of that nucleosome away from other heterochromatin. G. Chromocenters labeled in an NIH-3T3 cell based on DAPI content. H. H4G4D forms distinct punctate structures adjacent to chromocenters in NIH-3T3 (left) and C2C12 (right) cells. I. H4G4D, but not H4 (top row), disperses HP1a in NIH-3T3 chromocenters. Plot represents max-normalized fluorescence intensities of DAPI (blue), GFP (green) and HP1a (purple) along the dotted lines in the DAPI images. J. Schematic of RIME method. K. Shared and unique H4-GFP and G4D-GFP interactors measured using RIME, highlighting core nucleosome components (shared) as well as macroH2A.1 (H4 only) and some nuclear lamin components (G4D only). L. Quantification of signal in H3K9me3 CUT&Tag peaks called from H4 samples, showing a decrease in signal in the same regions in G4D cells. Peaks are ordered based on signal strength in the H4 samples.

To improve the resolution of this exclusion effect, we turned to a well-defined model of structural heterochromatin. Heterochromatin and euchromatin are characteristically compartmentalized into different parts of the nucleus through self-association of heterochromatin and exclusion of euchromatin [29]. A classic example of this process is chromocenters, structures composed of pericentric heterochromatin and found in mouse and *Drosophila* [30]. In this model of compartmentalization, an irreversibly “euchromatinized” nucleosome that promotes an open chromatin state should be partitioned away from dense heterochromatin (**Figure 2F**); indeed, recent work has suggested a model wherein compartmentalization behavior is encoded in individual nucleosomes [31]. To examine whether such compartmentalization occurs, we expressed the mutant H4G4D, which had displayed the largest extent of partitioning, in NIH-3T3 cells and C2C12 cells, both of which have large, easily visualized chromocenters (**Figure 2G**). Strikingly, H4G4D appeared as punctate structures adjacent to chromocenters in both cell types (**Figure 2H**). Although chromocenters seemed to be visually unchanged by the expression of G4D, the partitioning was found to also disrupt HP1a localization in chromocenters where G4D was not present (**Figure 2I, Figure S3C**), suggesting that although its effect on nucleosomal state occurs in *cis*, this can cause downstream *trans* effects via altered partitioning of chromatin components.

We reasoned that the effects observed above were mediated by primary effects in *cis,* wherein H4G4D either prompts the recruitment or prevents the binding of chromatin regulatory proteins. To this end, we performed rapid immunoprecipitation mass spectrometry of endogenous protein (RIME) [32] on the GFP tag in both H4 and H4G4D cell lines (n = 3 per cell line), allowing us to identify shared and unique interactors between the wild type sequence and the G4D mutant (**Figure 2J, Figure S3D-E**). Strikingly, although many nucleosomal components were shared between the two histones, H4G4D displayed a unique lack of interaction with macroH2A.1 (*p*=0.03), an established heterochromatin component (**Figure S3F**) [33], supporting our hypothesis that the mutation was creating heterochromatin-resistant nucleosomes in *cis* (**Figure 2K**). Interestingly, H4G4D also displayed interactions with prelamin A/C, lamin B1, and nucleophosmin (p < 0.05 for all, **Figure S3G**), suggesting a possible higher affinity for the nuclear membrane and nucleolus and suggestive of its disruption of H3K9me3 at these sites. Finally, quantifying signal strength at H3K9me3 peaks using CUT&Tag [34] confirmed a loss of signal in peaks identified in H4 cells (**Figure 2L**). Together, these results suggest a model in which certain mutations like H4G4D are able to disrupt the ability of their nucleosomes to be heterochromatinized, the effects of which extend to other nucleosomes and possibly larger bodies of heterochromatin as well.

### H4G4D alters the transcription factor binding chromatin landscape and functional TF-regulated outcomes

Having established a potential mechanism for H4 tail mutant activity, we next asked whether these altered chromatin states affect cellular responsiveness to transcription factors (TFs). In both natural and engineered systems, TF activity, templated on DNA-encoded binding sites and regulated by the epigenetic landscape and its plasticity, is a major driver of cell behavior [35–36]. We hypothesized that mutant-induced chromatin changes might reprogram the regulatory landscape, altering how cells interpret TF input and thereby modulating TF-driven phenotypes such as proliferation (**Figure 3A**). To explore this possibility, we first collected ATAC-seq data on several H4 tail mutants, focusing on motif-level changes in chromatin accessibility. A number of motifs associated with heterochromatin domains (H3K9me3), including a range of zinc finger proteins [37], showed elevated accessibility in the presence of H4 tail mutants including G4D (**Figure S4A-C**). These observations suggested that H4 mutants might unmask previously inaccessible binding sites by disrupting the heterochromatin that contains them. However, there are many factors affecting whether a TF with an accessible site truly binds: motif accessibility alone does not predict whether these TFs actually exert functional consequences [38–39].

**Figure 3:**
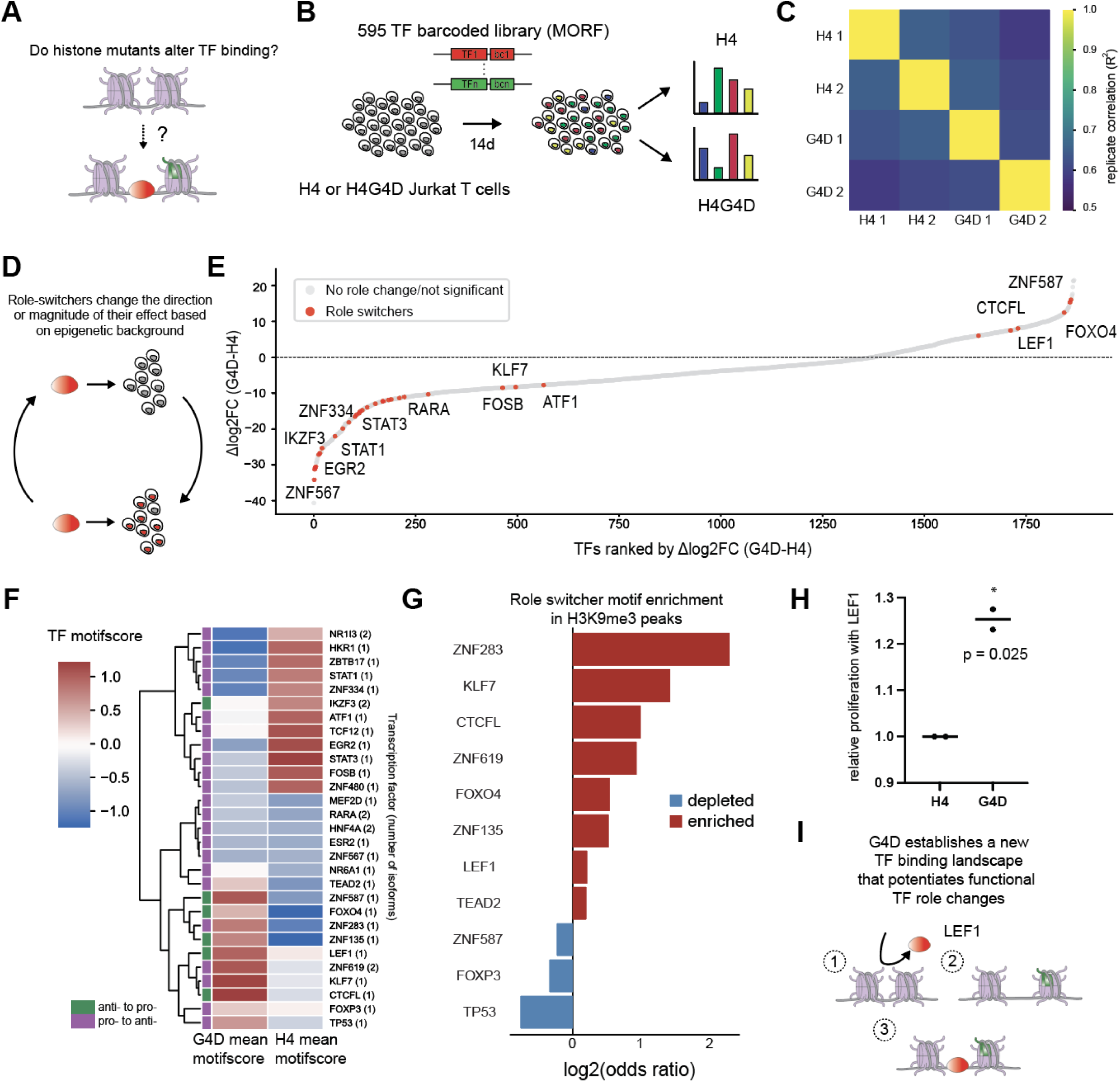
**H4 mutants alter the transcription factor binding landscape and functional TF-regulated outcomes.** A. We asked whether the incorporation of mutant histones (green) into native chromatin (pink) could remodel nucleosome arrangement and alter transcription factor (red) binding. B. Diagram showing layout of Jurkat MORF screen. Jurkat T cell lines were generated, infected with the lentiviral MORF library, and samples were collected at 14 days. Barcode abundance was measured to compare relative proliferation in different epigenetic backgrounds. C. Correlation matrix between replicates of MORF595 H4 and G4D screens. D. Schematic of role-switcher concept. E. Ranked plot of differential in changes in barcode abundance between t0 and t14 between G4D and WT cells. Cells denoted as “role switchers” denote padj < 0.05 in either condition and change in log2FC sign between WT and G4D. F. ATAC-seq motif scores for role-switchers (motifs only used when available) in WT and G4D cells, showing changes in motif accessibility in a number of anti-proliferative to pro-proliferative role switchers (green). G. Enrichment of role-switcher motifs in annotated Jurkat H3K9me3 peaks over background, showing enrichment of many role-switchers in heterochromatin. H. LEF1 increases proliferation in G4D cells. I. Our proposed paradigm is that G4D expression (green histone) alters the chromatin landscape and allows TFs (such as LEF1 in red) to bind.

To directly assess TF activity in mutant-expressing cells in a therapeutically relevant context, we generated stably expressing H4 and H4G4D Jurkat T-cells, a context where epigenetic barriers often limit TF-driven reprogramming [40]. After validating decrease of H3K9me3 upon G4D expression (**Figure S4D),** we performed high-throughput TF proliferation screens in replicate (R^2^ = 0.62-0.65 between screen replicates) using the MORF595 library: a barcoded pool of 595 human transcription factors and controls [41] (**Figure 3B-3C**). We also collected ATAC-seq data [42] on histone-expressing (but non-MORF library expressing) cells. We reasoned that comparing proliferation in different epigenetic backgrounds could highlight epigenome-specific potentiation or blocking of particular TF effects, which may cause particular TFs to “switch” the direction of their effect on proliferation, or transition from having no significant effect to having an effect (**Figure 3D**). Strikingly, several TFs, including the Lymphoid enhancer-binding factor 1 (LEF1), previously found to regulate T cell maturation and proliferation [43–44], appeared as “role switchers” in our screen (**Figure S4E, Figure 3E**). ATAC-seq data from WT and G4D cells revealed that LEF1 and other such “role switchers” displayed increased motif accessibility in the G4D epigenetic background (**Figure 3F**), and were enriched in H3K9me3 regions (**Figure 3G**), consistent with a model in which disruption of the heterochromatin landscape increases motif accessibility for certain TFs. The licensing of LEF1 to promote proliferation was validated in WT and G4D cells expressing LEF1 (**Figure 3H**). Together, these results present a proof of concept: histone mutants like G4D, through restructuring the chromatin landscape, can potentiate cellular plasticity to overexpressed TFs and result in functional TF activity and downstream cellular outcomes, similar to the effects of histone variants in nature (**Figure 3I**).

### Evolved and designed histones alter the transcription factor binding landscape

Up until this point, our efforts had been focused on single histone mutations. We next asked whether we could use these insights to combine histone mutations in a rational manner, to enable additional tunability of the chromatin landscape. Our initial efforts to express H4 proteins with multiple histFlow-nominated mutations in the globular domain often failed to result in incorporation (**Figure S5A**), while the tail was able to sustain many mutations and still allow integration into chromatin (**Figure S5B**). Given its small size relative to the whole histone, as well as its *cis*-acting behavior, we next asked whether mutated H4 tails alone were capable of chromatin remodeling when tethered to a DNA-binding domain (DBD) and recruited to a regulatory region (the HT-Recruit system, which has been demonstrated to be sensitive to nucleosome formation at its promoter [45]). A near-saturation-level library of mutant H4 tails (**Figure S5C**) fused to the tetR DBD and recruited to either a low-expressing promoter (activation screen) or a strong promoter (repression screen) failed to cause any change in expression (**Figure S5D-E**), suggesting that H4 tails are likely insufficient to cause meaningful changes in chromatin structure when merely recruited to DNA, and instead require nucleosomal context to function. We concluded that the wildtype H4 globular domain could be kept constant and the tail modulated to access additional cellular functions, calling this design *synH4* (**Figure 4A**).

**Figure 4:**
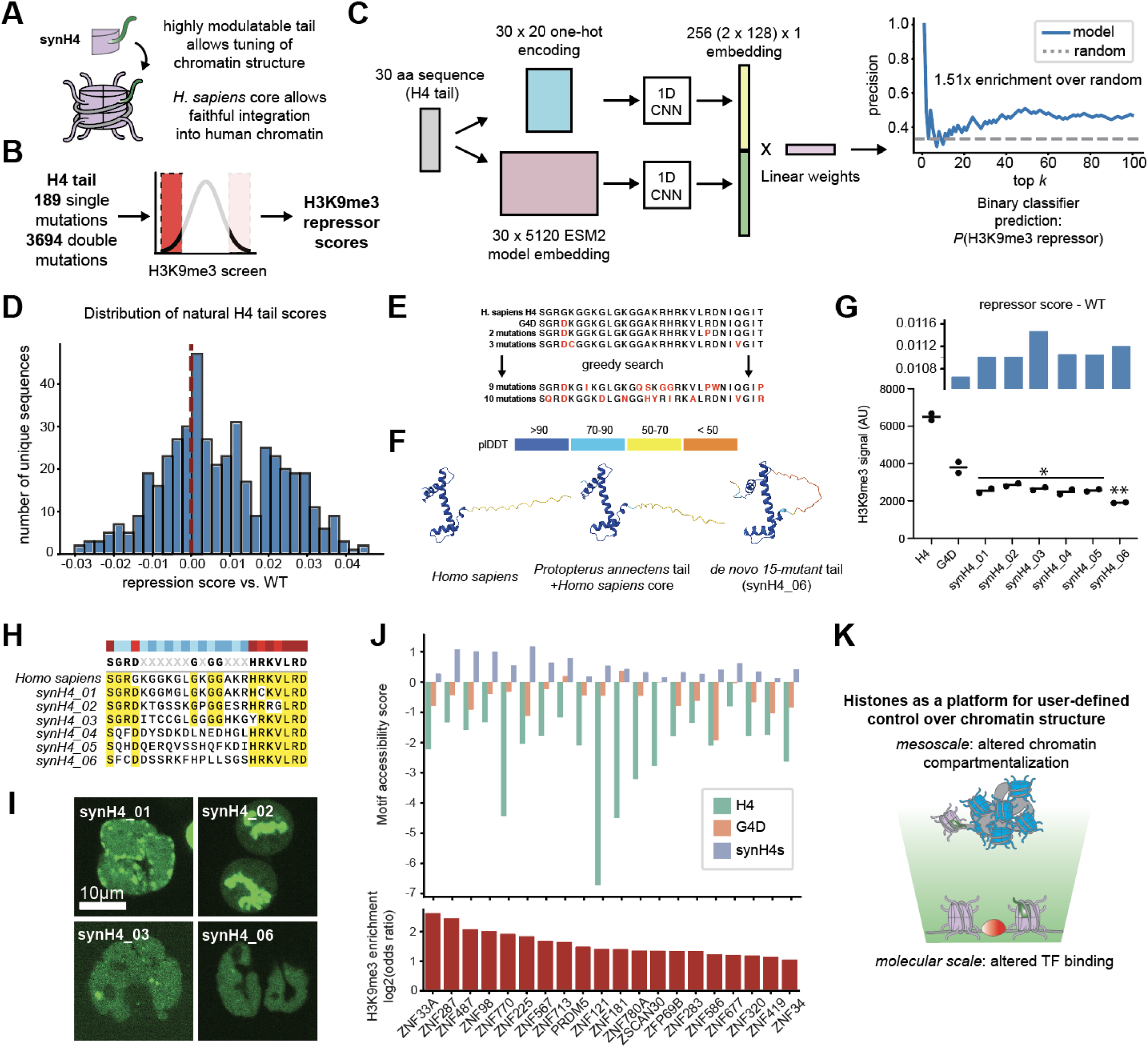
**Evolved and designed histones alter the transcription factor binding landscape.** A. Schematic of synH4 design, consisting of a wildtype (*H. sapiens*) chassis (pink) and a modular tail (green). B. Double-mutant H3K9me3 screens were conducted on a library of 189 single mutations and 3694 double mutations. C. Schematic of histForge training: H4 tail sequences are used to generate both ESM2 embeddings and one-hot encodings, then processed in parallel by 1D convolutional branches, and finally passed to a fully connected classifier. The final output was a repression probability between 0 and 1 (precision@N chart shown, for the top ranked (most H3K9me3 repressive) sequences). D. Distribution of “natural” H4 tails from Uniref, showing a spread of predicted repressor activity around the WT (dotted line). E. Diagram of greedy search algorithm for nominating synthetic tails: mutations were added that were predicted to increase H3K9me3 repressor score. F. AlphaFold-predicted structures of both *Protopterus annectens* and *de novo* synH4s compared to *Homo sapiens* H4, showing little disruption to predicted structure aside from small changes in predicted order at the N-terminal tail in synH4_06. G. Measured decreases in H3K9me3 in cells stably expressing nominated synH4s, with predicted repressor scores from histForge plotted above. * denotes p < 0.1 and ** denotes p < 0.05. H. Alignment of *Homo sapiens* H4 tail with the tails of synH4s 01-06. I. Proper incorporation of synH4s into human chromatin, including mitotic chromosomes (top right, synH4_02). J. Accessibility increases (top histogram) in ATAC data collected from synH4-expressing cell and corresponding H3K9me3 motif enrichment (bottom histogram), showing changes in H3K9me3-enriched motifs. K. synHistones allow for engineered changes at both the molecular scale (bottom) and mesoscale (top).

We then returned to our original question of whether combinatorial mutations could result in diversified function. To address this question, we constructed an “epistatic variant” histFlow library where individual tail mutants chosen from the results of our previous H4 screens, as well as a subset of all possible double mutants from that list, could be interrogated to determine their effects on H3K9me3 (**Figure 4B**), and performed H3K9me3 screens as described previously (**Figure S5F**). Having collected this new dataset, we next asked whether our combinatorial histFlow data could be used to build a predictive model of histone tail behavior that could assist in synH4 design. We trained a convolutional neural net on combined one-hot encodings [46] and ESM2 embeddings [47] of all tail sequences used in the double-mutant histFlow screen, and trained a binary classifier to predict the probability that a given sequence was a H3K9me3 repressor. Our model, which we called histForge, was able to predict the highest-ranking H3K9me3 repressors at an enrichment of 1.51x over random guessing (**Figure 4C**). We used histForge to generate repression scores for all unique eukaryotic H4 tail sequences, revealing a range of predicted repression activity compared to the *Homo sapiens* tail (dotted line) (**Figure 4D**). Our model predicted a range of behavior, with many sequences both more and less likely to be heterochromatin repressors, suggestive of the higher sequence diversity in the H4 tail region that we noticed while constructing our original mutant library. Applying this training scheme to a corresponding ATAC-see screen, conducted with the same library, resulted in appreciable albeit less accurate classification (1.17x over random, **Figure S5G**), nevertheless suggesting the generalizability of this approach.

Finally, we used histForge to assist in the design of fully synthetic histones. Starting with the G4D sequence as an initial “anchor” state and adding mutations, we implemented an iterative greedy search algorithm that would maximize the predicted total effect on H3K9me3 (**Figure 4E**). We also nominated a naturally occurring H4 tail, from the West African lungfish *Protopterus annectens*, that was scored highly by our histForge predictor and contains the G4D mutation. Predicted folds for nominated synH4s suggested high similarity to human H4, barring some changes in predicted order at the tail region (**Figure 4F**). Designed synH4s successfully decreased H3K9me3 levels past the level of G4D alone in K562s (**Figure 4G**) as well as in Jurkat T cells (**Figure S5H**). Moreover, in support of our earlier findings that the H4 tail was highly amenable to mutation, despite their significant divergence from the wildtype sequence (up to 62.5% difference by sequence composition, **Figure 4H**), synH4s integrated correctly into interphase and mitotic chromatin (**Figure 4I**). ATAC-seq profiling of synH4-expressing K562s demonstrated an increase in motif accessibility at several ZNF binding sites enriched in H3K9me3 regions (**Figure 4J**). Importantly, these sites displayed increased motif accessibility beyond the effects of G4D alone; since these are bulk accessibility measurements, this suggests that synH4s have a greater probability than G4D of disrupting heterochromatin at any given site, leading to an overall decrease in population-level H3K9me3 as well as bulk-measurement increases in ZNF accessibility.

Together, these results establish a chassis-tail paradigm (synHistones) for programmable chromatin: the histone fold provides a minimal, functional scaffold deposited by endogenous H4 chaperones, while the attached tail serves as a tunable interface for chromatin signaling. This system opens new avenues for synthetic epigenetics, enabling rational design of nucleosome-like particles that modulate chromatin state with modular precision across relevant spatial scales (**Figure 4K**).

## Discussion

Here, we present a unified framework, spanning high-throughput screening (histFlow), imaging, genomics, and synthetic design (histForge), that uncovers how amino-acid-level perturbations to histone sequence can reshape chromatin architecture in human cells and positions histones as highly engineerable substrates. We identify histone variants capable of disrupting chromatin compartmentalization and reprogramming the transcription factor binding landscape through both *cis* and *trans* mechanisms. A key takeaway from our work is that the mutation-to-function landscape is highly complex in histone proteins: many mutations do not “break” the protein but rather tune its interaction with the local chromatin environment, while maintaining its basic properties of incorporation and stability. Indeed, all screened amino acid mutations were mined from evolutionary histone variants and did not affect viability in human cells, supporting their compatibility with life. This suggests a histone evolutionary landscape where mutations are not all neutral or dominant negative, but have gain-of-function effects that may be tuned in various organisms by factors such as genome structure, environment, or developmental dynamics [48]. Similar concepts have been explored in prior work; for example, in understanding the evolution of DNA damage repair activity in rodents with diverse lifespans [49]. It will be interesting to see if this holds true for other chromatin proteins in future work.

A second key finding of our work is that particular single point mutations within the H4 tail, a region previously considered secondary to DNA-contacting domains, can dominantly decompact heterochromatin and redistribute key chromatin regulators such as HP1a and macroH2A. We chose to deeply characterize one such mutation, H4G4D, finding that it drives the formation of puncta excluded from chromocenters and, through its effect on heterochromatin, licenses TF access to otherwise inaccessible motifs. This altered chromatin environment enables previously silent TFs to engage the genome and drive proliferative outcomes, revealing that an epigenetic barrier can be abolished by histone expression alone. The H4 tail has been previously shown to interact with pioneer factor TFs [25], supporting an important role for its physical effect on nucleosomal organization. Although the H4 tail is disordered and therefore difficult to structurally characterize, our findings, especially the fact that G4D and other H4 tail mutations are not always on PTM sites, bolster a model in which both intrinsic physical properties and PTMs converge to regulate histone tail function, and alterations in the extent of intrinsic disorder, such as those seen in synthetically designed histones, may offer future insight as to the role of tail conformational diversity in nucleosomal organization.

Finally, through the integration of combinatorial mutational scanning and protein language models, we further demonstrate the potential to design synthetic histones with predictable effects on heterochromatin structure. To this end, we developed histForge as a framework for sequence-based prediction and rational design of histone variants that modulate chromatin state. By integrating screening data on multiple-mutant histones with protein language models, histForge enables prediction of functional histone sequences across an expansive mutational space. Using this approach, we identified synthetic histones that outperform natural variants in heterochromatin repression, despite the combinatorial complexity of the sequence landscape (∼10^39^ possible combinations). We expect that screening across larger and more comprehensive libraries for additional chromatin phenotypes will only strengthen the predictive ability of such models, positioning our approach as a generalizable platform for *in silico* design of chromatin-regulatory proteins.

Together, our results establish evolutionary-scale screens as a robust method to explore chromatin biology, and establish histones as a new substrate for synthetic biology. Just as synthetic transcription factors and CRISPR effectors have transformed gene control, engineered histones offer a route to modulate the physical state of chromatin itself. Unlike most chromatin-modifying tools, which act indirectly or transiently, histone variants become embedded in the chromatin fiber, positioning them as durable, inheritable actuators of nuclear organization. Their transgenic nature also suggests synH4s could be integrated into synthetic gene circuits [50], allowing for increasingly complex and time-variant chromatin programming. Moving forward, we envision histone design being applied across diverse contexts: to override differentiation barriers in cell therapy, to encode self-limiting gene expression circuits, to reprogram TF activity in exhausted or senescent cells, or to create orthogonal chromatin compartments for synthetic genome engineering.

## Acknowledgments

We are grateful to the Buenrostro Lab for helpful feedback throughout this body of work, as well as for helpful discussions with N. Nagaraj. We would like to thank the Harvard Bauer Flow Cytometry Core (J. Nelson) for assistance with flow and the Harvard Mass Spectrometry Core (M. Chen and S. Kolakowski) for assistance with RIME mass spectrometry. We also gratefully acknowledge the use of the Revvity Opera Phenix High-Content/High-Throughput imaging system at the Broad Institute, funded by the S10 Grant NIH OD-026839-01.

## Author contributions

S.G.J. and J.D.B. conceived of the project and designed experiments. S.G.J. performed all cloning, screens and flow validation, ATAC-seq experiments, confocal imaging experiments, data analysis, and computational modeling. S.N. performed RIME sample preparation. S.G.J. and A.E. performed CUT&Tag experiments. S.G.J. generated all cell lines with assistance from W.G. and A.D-C. M.Q. assisted with high-throughput imaging. E.O. and M.Q. assisted with MORF595 library generation. M.A.H. assisted in data interpretation and visualization.

R.Z. assisted with ESM2 embedding generation. J.D.B. supervised all aspects of the study. S.G.J. and J.D.B. wrote the manuscript with input from all authors.

## Data and materials availability

As of 9/12/2025, we are in the process of uploading processed data files to a GSE accession and code to GitHub and Zenodo. This preprint will soon be updated accordingly.

## Materials and Methods

### Cell culture

K562 cells (ATCC CCL-243) and Jurkat cells (ATCC TIB-152) were cultured in RPMI-1640 medium supplemented with 10% fetal bovine serum (FBS) and 1% penicillin-streptomycin at 37°C in a humidified incubator containing 5% CO₂. NIH-3T3 cells (ATCC CRL-1658) and C2C12 cells (ATCC CRL-1772) were cultured in DMEM supplemented with 10% fetal bovine serum (FBS) and 1% penicillin-streptomycin at 37°C in a humidified incubator containing 5% CO₂. To prevent FBS-dependent changes in proliferation and cell growth, multiple FBS lots were tested and a lot that resulted in consistent and robust growth in multiple cell lines (K562, LentiX) was selected and used for all experiments. For doxycycline (dox) induction systems, doxycycline powder was dissolved in water to make a 1 mg/mL stock solution, which was aliquoted to minimize freeze-thaw cycles and kept at-20°C until use. Dox was added to cells to a final concentration of 1 ug/mL, and this concentration was used for all experiments involving dox. Histone expressing cells were selected with geneticin/G418 (800 µg/mL, diluted from a stock solution of 50 mg/mL) for 8 days. HT-recruit cells were selected using puromycin at a final concentration of 2 ug/mL, from a stock solution of 10 mg/mL, for 5 days. In both cases, the effectiveness of selection was tested by treating a subsample of cells with dox and ensuring the % GFP positive was ∼100%, measured through flow cytometry.

### Computational nomination of histone sequence variants

To generate a database of histone sequence variants from nature, protein sequences from Uniprot queries (i.e. “histone h4”) for each of the core histones (H2A, H2B, H3, and H4) were downloaded. Protein sequences were current to the Uniprot database as of 11/1/2023. The resulting sequences were filtered to 1. Remove sequences annotated as fragments, 2. Remove any possible non-histone enzymes and chaperones (i.e. “Histone H3 acetyltransferase”, “Histone H4 chaperone”, “Histone H3 methyltransferase”), and finally to 3. Truncate sequences to those that were the same length as the human protein. The resulting sequences were aligned using the Hidden Markov Model (HMM) tool jackHMMer. The resulting alignment was used to enumerate the number of total differences between the *H. sapiens* sequence (Uniprot accession #s P62805, P33778, Q6FI13, and Q6NXT2) and the query sequence, for each sequence in the subset. Query sequences that were more than 20% different by amino acid content were discarded, and the resulting list of single amino acid changes was used to build the library for that histone. Additionally, alanine mutants for each position in the protein, as well as arginine, glutamate and methionine mutants for each lysine were added to its library. Phylogenetic trees for each histone family were generated using the Phylo module in biopython (https://biopython.org/wiki/Phylo), and alignments for each histone family as input. All code used to generate phylogenetic trees is provided.

### Double-mutant library generation

A collection of the top tail mutations (by signed FDR) from ATAC and H3K9me3 screens in H4s was compiled, and a subset of all possible 2-mutant combinations from this set was generated with custom MATLAB code and sampled in a Twist library and cloned into the H4 inducible backbone (see Cloning). 189 single mutations and 3694 double mutations were accurately called from the resulting library.

### rTetR-tail library generation

A Twist library encoding every possible amino acid mutation in the H4 tail was synthesized and designed to be cloned upstream of the rTetR construct in the HT-recruit construct.

### Cloning

For each histone pool, a human codon-optimized copy of the wild-type histone sequence (Uniprot accession #s P62805, P33778, Q6FI13, and Q6NXT2) was cloned into the pCW57-RFP-P2A-MCS (Neo) vector, which was a gift from Adam Karpf (Addgene plasmid # 89182). Sequences were generated with no introns. This construct allowed for inducible expression of the histone gene in combination with turbo RFP (tRFP) using the P2A self-cleaving peptide, and contains the rTetR gene for all-in-one induction of the histone protein.

**Figure 1:**
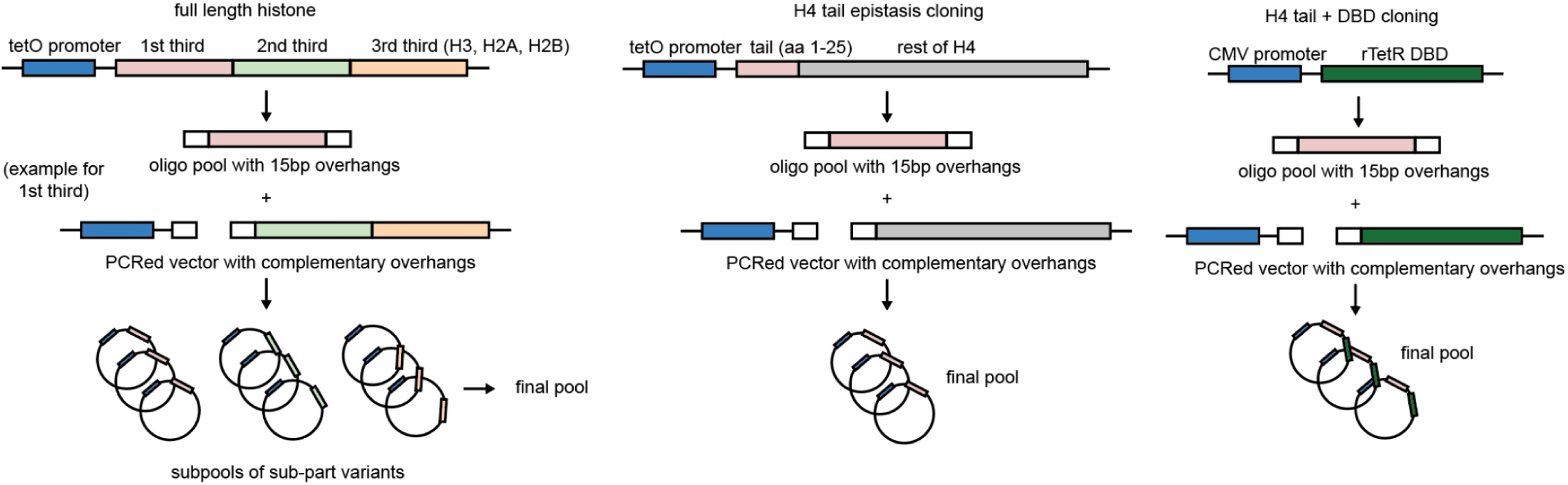

For each histone, the total length of the protein was split into either two or three equal-sized parts. For each section, a library of oligonucleotides containing the relevant mutations and 3’ and 5’ overhangs of 15 bp was synthesized using the Twist oligonucleotide synthesis service, amplified with primers complementary to the 15 bp overhangs, and ligated into a vector generated by amplification with primers complementary to the 15bp overhangs. Ligations were performed with Gibson HiFi Assembly, with a ratio of 9:1 fragment to vector, and the resulting ligation product was transformed into ElectroMAX competent cells by electroporation as per the manufacturer’s protocol. Transformants were spread onto BioAssay carbenicillin plates and cultured at 30°C overnight, after which coverage was estimated using the colony count on a 1:1000 and 1:10000 dilution plate to confirm > 500x coverage. At this step, up to 10 colonies were picked and cultured for individual validation through plasmid sequencing (Plasmidsaurus) for verification, to ensure mutations were in the correct part of the histone and that no other mutations had occurred. Colonies were scraped and collected in 100 mL of media, cultured for an additional 3 hr, and plasmid DNA collected with a Qiagen MaxiPrep kit. Final pools were created by combining subpools in equal molar fractions. **Note:** the above approach was used for the original histone mutant libraries, the H4 tail epistasis library, and the tail+rTetR library. For single point mutations, mutations were encoded using site directed mutagenesis in the wildtype protein of interest using overlapping primers. NEB Stable cells were transformed with the resulting amplified vector, and colonies picked from carbenicillin plates containing the resulting transformants and plasmid sequenced using Plasmidsaurus for verification.

For synH4 constructs, mutations were encoded in two complementary oligos with overhangs complementary to the 5’ UTR of the H4 sequence and the globular domain. Oligos were used as primers for long PCRs, annealed, and transformed into competent cells as described above. Sequences were confirmed using Plasmidsaurus sequencing.

### MORF595 library cloning

The MORF595 library was cloned by PCR amplifying TFs with V5 fusion out of the original MORF library (Joung et. al. *Cell* 2024) and Gibson assembly into the pLX307 plasmid. In brief, 595 TF ORFs with V5 fusion in the MORF library were PCR amplified with a forward primer specific to ORFs with V5 fusion for 6 cycles total.

MORF_595_Forward: GTGAGGCTAGCATCGATTGATCAACAAGTTTGTAC

MORF_595_Reverse: TTGTAATCCAGAGGTTGATTGTCGACTTAACGC

The original plasmids were digested with DpnI restriction enzyme (NEB) for 2 hours at 37C. PCR amplicons were then PCR purified using Qiagen MinElute PCR Purification kit. pLX307 plasmid was restriction enzyme digested using MluI and ClaI restriction enzymes (NEB) and the digested plasmid was agarose gel purified using a Zymoclean Gel DNA recovery kit. Plasmid library was ligated with NEBuilder Gibson Assembly and electroporated into Stbl4 electrocompetent bacteria with final representation >500x per ORF sequence.

### Cell line generation

Lentiviral vectors encoding histone mutant libraries were produced in HEK293T (LentiX) cells using standard packaging protocols. Packaging plasmids used were VSVG and the 2nd generation lentiviral packaging plasmid psPAX2, and lentivirus was harvested either 2 days post-transfection, or both 2 and 3 days post-transfection for increased yield (with fresh media replacement after the 2-day timepoint collection). For screens, K562 cells were transduced with the lentiviral particles at a multiplicity of infection (MOI) of ∼0.3 to ensure single-copy integration. For non-screen applications (i.e. validation of single mutant cell lines), MOI did not surpass 0.3 (evidenced by % GFP expression after a test doxycycline treatment after transduction). Transduced cells were selected using geneticin (800 µg/mL) for 8 days prior to downstream applications. For all screens, cells were maintained at a representation of at least 2000x per replicate per screen in 75 mL flasks.

### Flow cytometry-based screening and validation (histFlow)

For epitope and ATAC screens, cells were grown in the absence of dox to 1000x coverage per replicate, before dox induction for 4 days. Each replicate was collected together and split into subpools for epitope staining (maintaining at least 1000x coverage per replicate). Cells for epitope screens were fixed with 1% paraformaldehyde for 10 minutes at room temperature, then permeabilized with 0.1% Triton X-100 for 10 minutes as per the Cell Signaling Technologies protocol. Cells were then blocked in PBS containing BSA (1%) and EDTA (10 mM) for 1 hr at 4°C, before washing in PBS and overnight incubation with one of the following primary antibodies: anti-H3K9me3 (Cell Signaling Technologies, #13969), anti-H3K27me3 (Cell Signaling Technology, #9733), and anti-H3K27ac (Active Motif, #39133), each diluted 1:500 in staining buffer (PBS with 0.01% Triton-X100, 1% BSA and 10mM EDTA). Following incubation, cells were washed 3x with PBS and incubated with Alexa Fluor 647-conjugated donkey anti-rabbit secondary antibody and DAPI.

For ATAC-see, cells were collected and treated with DNAse I in media with diluted DNAse I buffer for 1 hr at 37°C to remove dead cells, which would overtranspose and potentially reduce signal in other cells. Cells were washed 3x in PBS, then fixed for 10 min at RT with 1% FA, followed by 3x PBS washes. Cells were permeabilized for 10 minutes with ATAC transposition buffer (as per Buenrostro et al., 2013) for 10m at room temperature. Tn5 (SeqWell) was loaded with 5’ biotinylated adapters by incubating with adapters and glycerol for 30m at room temperature. Cells were transposed with the biotinylated transposome for 30 minutes at 37°C with shaking, followed by 3x spin-down and wash in Tn5 wash buffer (50mM EDTA, 0.01% SDS in 1X PBS). Cells were then spun them down one more time and left in staining buffer overnight to block. The next morning, cells were incubated in H3S10ph antibody (Sigma) for 1hr RT in staining buffer, spun down and washed 3x in PBS, and then incubated with Alexa Fluor 568-conjugated secondary antibody (goat anti-rabbit), Alexa 700-conjugated streptavidin, and DAPI for an hour at RT, followed by a final set of 3 washes. **Note:** This method (fixation, staining) for ATAC-see measurement was validated using a similar readout to that used by Ishii et. al., where signal inside and outside of NIH-3T3 chromocenters was quantified. The choice to stain for mitotic cells was taken from the same publication. FACS was performed using a BD FACSAria III. The top and bottom 10% of AF647 (epitope) or AF700 (ATAC) cells were collected after gating out mitotic cells in the ATAC screen (high 568) and debris.

Mutant validation was performed as described above for pooled libraries, with some slight changes. 1. Flow analysis was performed with an Attune CytPix Flow Cytometer since subsequent cell recovery was not needed. 2. Far-red secondary antibodies and streptavidin were changed to Brilliant Blue conjugated versions (400-450 nm) and DAPI staining was removed, to accommodate the Attune laser set. All staining and transposition was scaled down to the relevant cell count.

### Screen sequencing and data analysis

For all histone flow and proliferation screens, genomic DNA was extracted from sorted cell populations using the Monarch Genomic DNA Prep Kit (New England Biolabs). Histone sequences were amplified out of the genomic DNA using custom primers. Primers included a 8bp barcode in addition to the complementary sequence in order to multiplex samples on a single run. Library preparation was performed using the Oxford Nanopore Technologies (ONT) Ligation Sequencing Kit (SQK-LSK110), following the manufacturer’s protocol. Sequencing was conducted on a MinION device equipped with R10.4 flow cells. Basecalling was performed using Dorado (ONT) with default parameters. Custom scripts were used to demultiplex reads and identify and quantify histone mutant variants present in each sample, maintaining a stringent 0-error threshold for both demultiplexing and variant calling.

For MORF screens, genomic DNA was extracted as detailed above, and TF barcodes amplified using custom primers. Barcodes were sequenced using Illumina NGS and mapped to a dictionary of barcode-TF pairs. Statistics for all screens were calculated using DeSeq2 on raw count data, and applying a correction using the counts for the GFP and mCherry controls.

### Proteomics

RIME was performed as described in Mohammed et al. 2016 with the following modifications. Cells were resuspended in PBS and 1/10th volume Crosslinking Solution (11% formaldehyde, 100mM NaCl, 1mM EDTA, 50 mM HEPES) was added. Fixations were performed by rotating for 8 min at RT then quenched with 1/20th volume of 2.5M glycine for 5 min at RT. Fixed cells were then washed twice with PBS + 0.5% NP-40, including 1 mM PMSF in the second wash. Pellets were stored at-80C.

Pellets were lysed in LB1 and LB2 as described in the protocol before resuspension in Sonication Buffer (10 mM Tris pH 8, 1 mM EDTA, 0.1% SDS). Chromatin was sonicated on a Covaris S220 at 140W, 10% duty and 200 cycles/burst such that the majority of fragments were between 150 and 600 bp. Buffer was supplemented with NaCl, EGTA, Na-deoxycholate, N-lauroylsarcosine and Triton-X 100 to match concentrations of the published LB3 buffer. Lysates were clarified by centrifugation at 20,000xg for 10 min and the supernatant was added to Protein A Dynabeads (Life Tech, 10002D) bound with 10 ug of anti-GFP antibody (Abcam, ab290). Samples were rotated overnight at 4C. RIPA washes were performed as described in the protocol. AMBIC washes were increased to 5 total washes, transferring to new tubes after the third and fifth washes to minimize detergent and salt carryover. Protein-bound beads were then frozen at-20C.

### On-bead Enzymatic Digestion

Magnetic beads were resuspended in 100 mM TEAB with 0.1% RapiGest. Proteins were digested with a Trypsin/Lys-C mix while incubating overnight at 37 °C in a shaking thermomixer. After digestion, samples were placed on the magnetic rack to separate the beads and the supernatant was transferred to a new tube. The beads were washed with 100 mM TEAB, which was combined with the previous supernatant solution. Samples were acidified with dilute trifluoroacetic acid (TFA) to a pH < 3 and then incubated at 37° C for 45 minutes to precipitate the RapiGest. They were then centrifuged at 16,000 g for 10 minutes, and the supernatant was transferred to a new tube. The samples were cleaned up with Pierce Desalting Columns in preparation for LC-MS/MS analysis.

### LC-MS/MS Analysis

Samples were analyzed by Q Exactive HF-X High Resolution Orbitrap (Thermo Fisher, Waltham, MA) coupled with Ultimate 3000 nanoLC (Thermo Fischer, Waltham, MA) at Harvard Center for Mass Spectrometry. Peptides were first trapped on a trapping cartridge (300µm x 5mm PepMap™ Neo C18 Trap Cartridge, Thermo scientific) prior to separation on an analytical column (µPAC, C18 pillar surface, 50 cm bed, Thermo scientific). Peptides were separated using a 120-min gradient (from 1 – 45% acetonitrile with 0.1% formic acid) with a flow rate of 300 nL· min−1. The mass spectrometer operated in data-dependent mode for all analyses. Electrospray positive ionization was enabled with a voltage at 2.1 kV. A full scan ranging from 350 to 1400 m/z was performed with a mass resolution of 6.0×104 and AGC target set to 3e6. The top ten most intensive precursor ions from each scan were used for MS2 fragmentation with normalized collision energy of 28 at a mass resolution of 3.0×104 and AGC of 1e5.

### Data Processing

Raw data was submitted for analysis in Proteome Discoverer 3.1 software (Thermo Scientific). The MS/MS Data was searched against the UniProt reviewed Homo sapiens database along with known contaminants such as human keratins and common lab contaminants. Quantitative analysis between samples was performed by label-free quantitation (LFQ). Sequest HT searches were performed using the following guidelines: a 10 ppm MS tolerance and 0.02 Da MS/MS tolerance; Trypsin digestion with up to two missed cleavages; deamidation (+0.984 Da) of asparagine and glutamine and oxidation (+15.995 Da) of methionine set as variable modification; minimum required peptide length set to ≥ 6 amino acids. At least one unique peptide per protein group is required for identifying proteins. All MS2 spectra assignment FDR of 1% on both protein and peptide level was achieved by applying the target-decoy database search by Percolator.

### Confocal and high-content imaging

Imaging was conducted in 96-well glass bottomed plates. For K562 imaging, wells were coated with a solution of 0.01% poly-D-lysine in water for > 1hr, after which they were washed with water and transduced K562 cells were plated onto the surface of the well by centrifugation at 300xg. For adherent cell imaging (NIH-3T3, C2C12), wells were coated with fibronectin at a concentration of 2 ug/cm2, for > 1hr, after which they were washed with water and transduced adherent cells were plated onto the surface of the well by centrifugation at 300xg. Cells were fixed immediately following plating for K562 imaging; for adherent cells, cells were incubated in normal conditions for at least 6 hours before fixing and staining, in order to fully adhere.

For imaging studies, cells were fixed with 4% paraformaldehyde. Immunofluorescence staining was performed as described above for flow screens, without spinning cells during each wash. Confocal imaging was conducted using a Nikon Eclipse Ti2, and 60x oil objective imaging was used unless otherwise noted for all images. while high-content imaging was performed on the Opera Phenix High-Content Screening System (PerkinElmer) using a 63x water objective. Image analysis for both confocal and high-content imaging, including quantification of nuclear localization and intensity measurements, was carried out using custom ImageJ and MATLAB scripts. Briefly, nuclei were segmented using DAPI staining, resulting in individual masks, and per-pixel GFP and/or H3K9me3 intensity within the resulting mask was quantified and either the mean over all pixels in the mask or the correlation between H3K9me3 and GFP calculated for each mask.

### Genomics

#### ATAC-seq

Assay for Transposase-Accessible Chromatin using Sequencing (ATAC-seq) was performed as previously described (Buenrostro et al., 2013) and performed identically for all cell types described in this paper. Briefly, cells were harvested during exponential growth and counted using an automatic hemocytometer. For each reaction, 10k viable cells were pelleted at 500 × g for 5 minutes at 4°C. Cells were washed once with 50 µL of cold 1× PBS and then resuspended in 5 uL, then permeabilized in 45 uL of transposition buffer for 10 minutes on ice. 2.5 µL Tn5 transposase (SeqWell) was added and the tagmentation reaction was incubated at 37°C for 30 minutes in a thermomixer at 300 rpm.

Following tagmentation, DNA was purified using a Zymo PCR Purification Kit (Zymo) and eluted in 10 µL of elution buffer. PCR amplification was performed using NEBNext High-Fidelity 2× PCR Master Mix with custom Illumina indexing primers, and the optimal number of cycles was determined using a qPCR side reaction (to avoid overamplification). Final libraries were purified using a Zymo PCR Purification Kit and eluted in 20 µL of elution buffer.

#### Cut&Tag

Cleavage Under Targets and Tagmentation (Cut&Tag) was performed using the EpiCypher CUTANA™ Cut&Tag Assay Kit. 100k K562 or NIH-3T3 cells per condition were washed in Wash Buffer (20 mM HEPES pH 7.5, 150 mM NaCl, 0.5 mM spermidine, protease inhibitor cocktail) and bound to Concanavalin A-coated magnetic beads for 10 minutes at room temperature with gentle rotation. Bead-bound cells were incubated with primary antibody (e.g., anti-H3K27ac, anti-H3K9me3, 1:100) diluted in Dig-Wash Buffer (Wash Buffer + 0.05% digitonin) overnight at 4°C with rotation.

Following primary antibody incubation, beads were washed and incubated with a secondary antibody (e.g., Guinea Pig anti-Rabbit IgG, 1:100) in Digitonin Wash Buffer for 1 hour at room temperature. After washing, the samples were incubated with pre-loaded pAG-Tn5 transposome (Epicypher) for 1 hour at room temperature. After washing, tagmentation was initiated by adding Tagmentation Buffer (10 mM MgCl₂ in Digitonin Wash Buffer) and incubating at 37°C for 1 hour. The reaction was stopped with 10 mM EDTA and 0.1% SDS and incubated at 55°C for 1 hour. DNA was purified using AMPure beads (1.5× ratio) and eluted in 20 µL of EB. Libraries were PCR-amplified using indexed primers and NEBNext Master Mix with 12–15 cycles, depending on yield and monitored using qPCR to prevent overamplification.

#### Genomics sequencing and data processing

Libraries were quantified using a Qubit Fluorometer and quality-checked on a Bioanalyzer (Agilent). Pooled libraries were sequenced on an Illumina NovaSeq or NextSeq using paired-end 50 bp reads. Reads were trimmed using `fastp`, aligned to the human genome (hg38, for K562 and Jurkats) or mouse genome (mm10, for NIH-3T3s) with `bowtie2`, and filtered to remove duplicates, mitochondrial reads, and low-quality alignments. Peaks were called using MACS2 with default parameters. Signal tracks were generated in bigWig format for visualization using `deeptools`. For analysis of motif coverage in peaks (ATAC and CUT&Tag), peaks (either generated from MACS2 or downloaded from GSE accessions/ENCODE) were analyzed for motif enrichment using chromVAR. For comparative enrichment of TF motifs in H3K9me3 regions, a GC-matched background peakset was generated based on the published ENCODE H3K9me3 peakset (ENCSR000APE) for K562s or GSE162605 for Jurkats, and enrichment in the corresponding H3K9me3 peakset vs the background set was calculated. For CUT&Tag analysis in mouse major satellites, enrichment of reads aligning to the dimerized mouse major satellite consensus (GenBank: V00846.1) was calculated.

### histForge modeling

Double-mutant tail screen DeSeq2 analysis was performed and signed FDR-adjusted p values generated for every single and double mutant in the screen. Each H4 tail sequence was encoded in two complementary ways: (1) a 30×5120 ESM2 per-residue embedding (`esm2_t48_15B_UR50D` model) generated using the `esm` package, and (2) a 30×20 one-hot encoding of the primary amino acid sequence. These two representations were processed in parallel by 1D convolutional branches (128 hidden channels, kernel size 5, adaptive max pooling), concatenated, and passed to a fully connected classifier. The final output was a repression probability between 0 and 1. To encourage biologically realistic predictions, we anchored the model to wild-type histone behavior. The wild-type H4 tail sequence was labeled as non-repressive and included with increased sample weight and repetition during training. Model training was performed using PyTorch with binary cross-entropy loss (BCEWithLogitsLoss, `reduction=’none’`), class-balancing using `pos_weight`, and per-sample reweighting to emphasize the wild-type anchor. Models were trained with Adam (learning rate = 1e-3) for 50 epochs using 80:20 train/test splits. We evaluated performance using AUC and precision@100 on held-out test data. Wild-type repression scores were tracked over training to confirm anchoring, and a histogram of predicted repression values was centered on the WT score to visualize enrichment or depletion across sequence space. Predictions on synthetic histones were reported as both raw and ΔWT scores.

## Supplemental Figures

**Figure S1:**
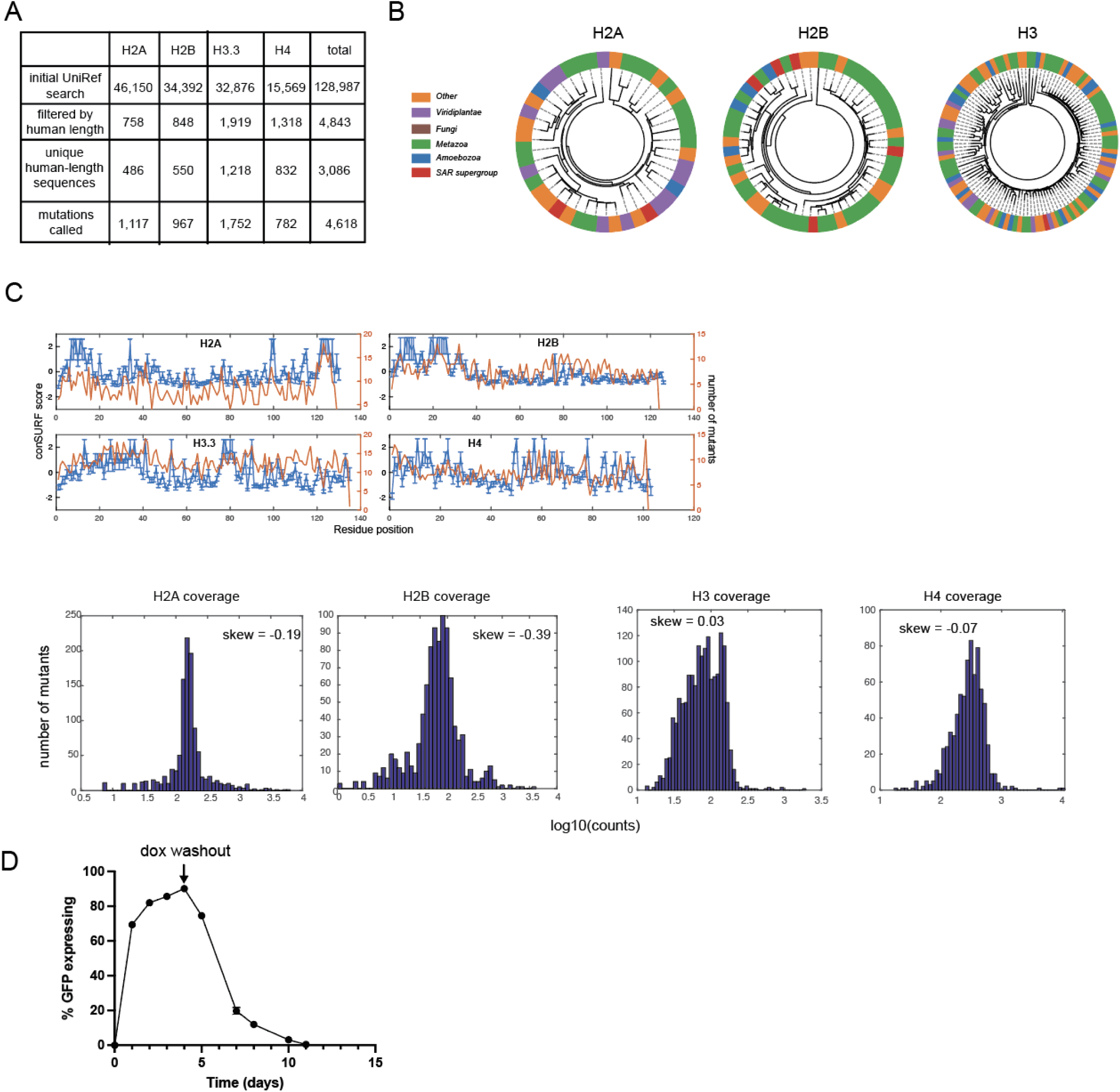
**A**. Breakdown of mutation calling steps for library creation. Histone proteins were called with UniRef, then filtered to human-length proteins, aligned, and mutations called. **B.** Phylogenetic trees for H2A, H2B, H3, constructed in the same way as H4 (Figure 1). **C. (top)** Number of mutants (orange) plotted over ConSurf conservation score (blue) **(bottom)** distribution of mutant representation in final libraries, with calculated skewness. **D.** Dynamics of induction and subsequent degradation of H4-GFP upon dox washout.

**Figure S2:**
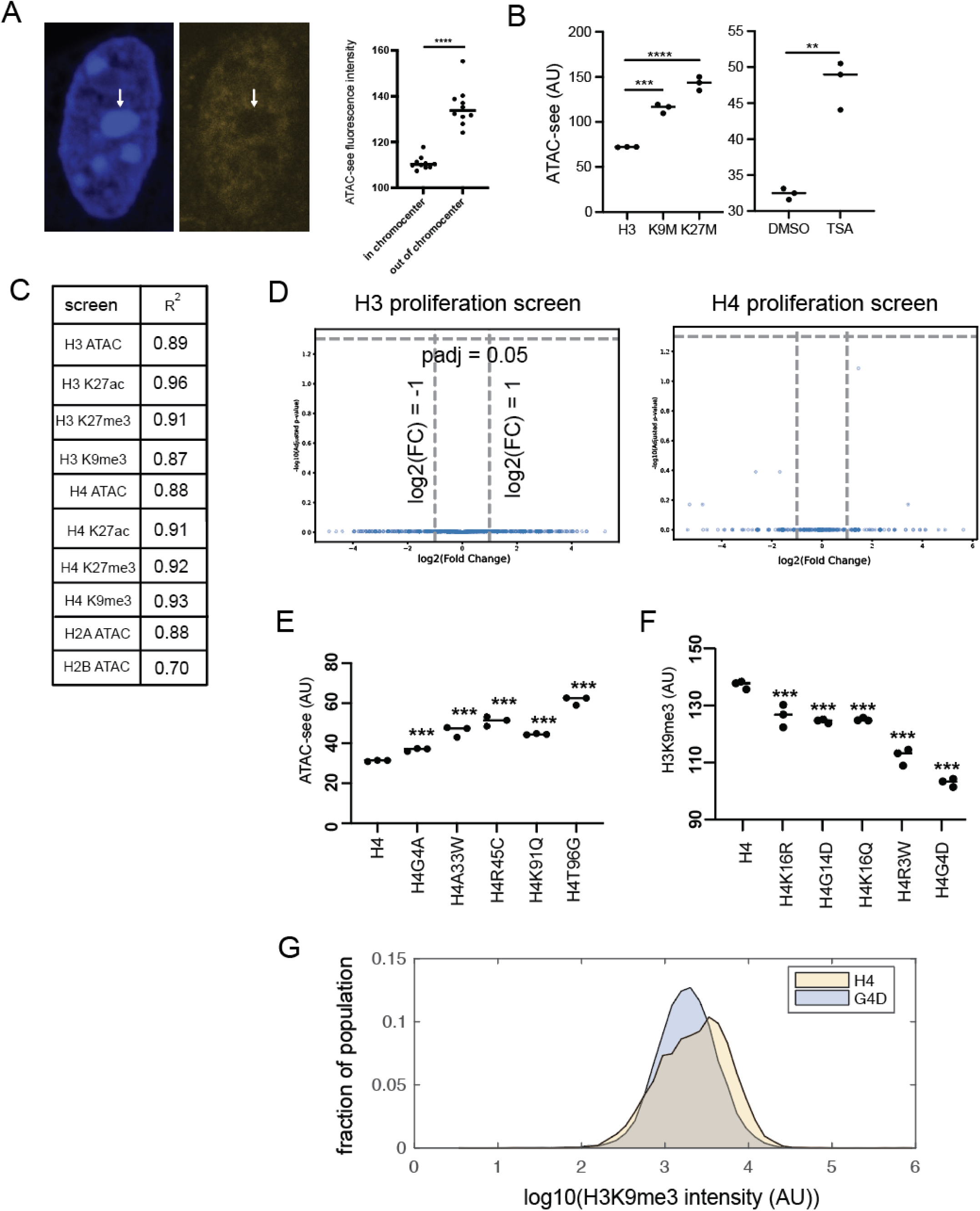
**A**. NIH-3T3 chromocenters showing high DAPI content (left) and low ATAC-see content (right), quantified on far right. **B.** H3 mutants and drugs increase accessibility of cells as measured through ATAC-see. **C.** Replicate correlation of screens in this study. **D.** H3 and H4 proliferation screens, showing no significant effect on proliferation at the time of cell collection. **E.** Validation of selected ATAC-see hits. **F.** Validation of selected H3K9me3 hits. **G.** Example of flow data comparing G4D to H4, showing decrease (∼30%) in mean H3K9me3 levels.

**Figure S3:**
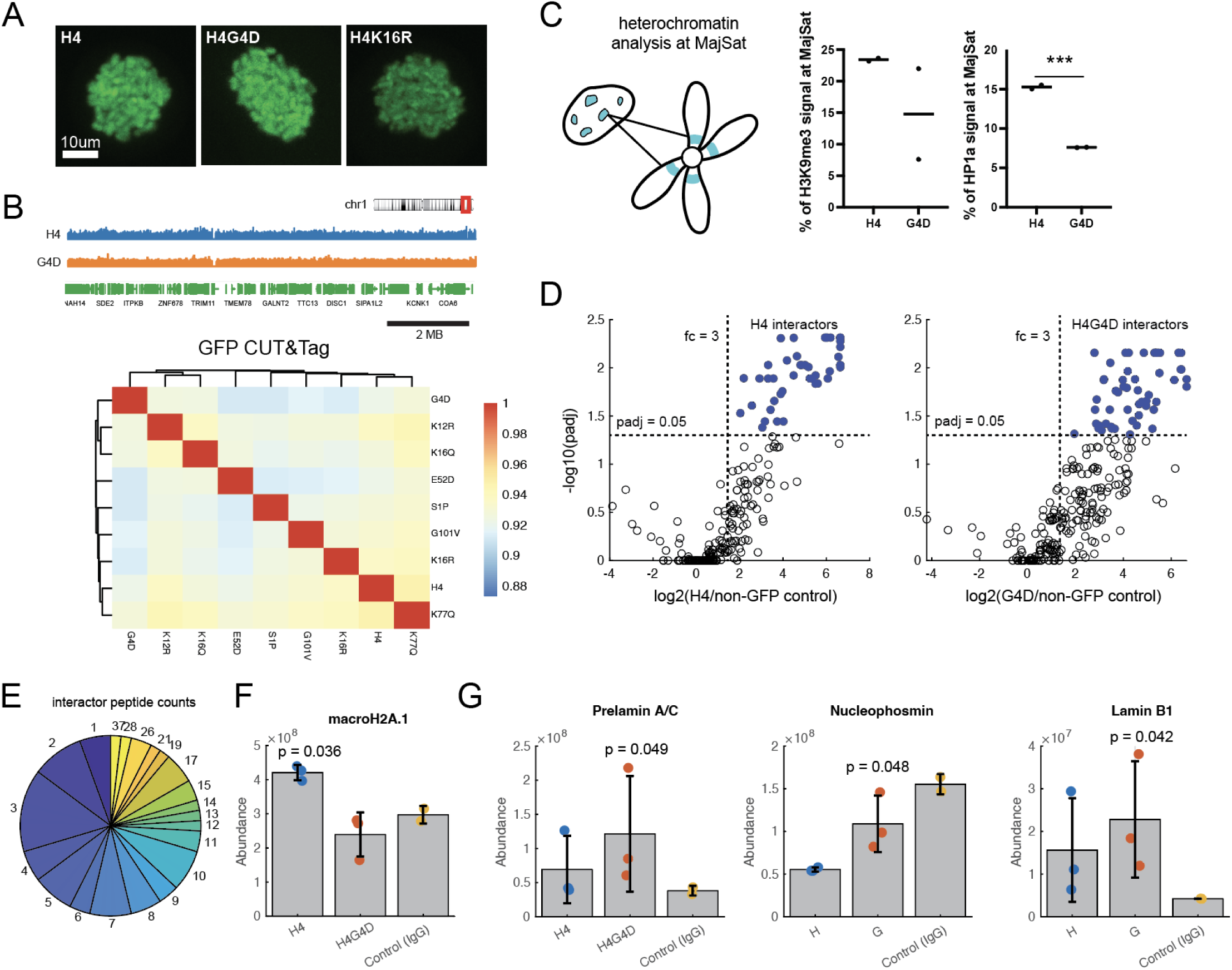
**A**. Mitotic chromatin demonstrates proper incorporation of mutant H4s. **B.** CUT&Tag of GFP-tagged histones with a range of mutations demonstrate high correlation (>0.9) between genomic enrichment levels across 50-kb bins. Sample tracks for H4 and G4D shown for a randomly selected region of chr1. **C.** CUT&Tag of H3K9me3 and HP1a in Major Satellite repeats in NIH-3T3s demonstrates a possible but not significant decrease in H3K9me3 (left) and a significant decrease of HP1a at MajSat sites (right). *** denotes p < 0.0005. **D.** Defining interactors of H4-GFP and H4G4D-GFP using RIME, by adjusted p-value cutoff and foldchange. **E.** Representative peptides per interactor, with most interactors represented by > 2 unique peptides. **F-G** Protein abundance for macroH2A.1, Prelamin A/C, Nucleophosmin, and Lamin B1, measured using RIME.

**Figure S4:**
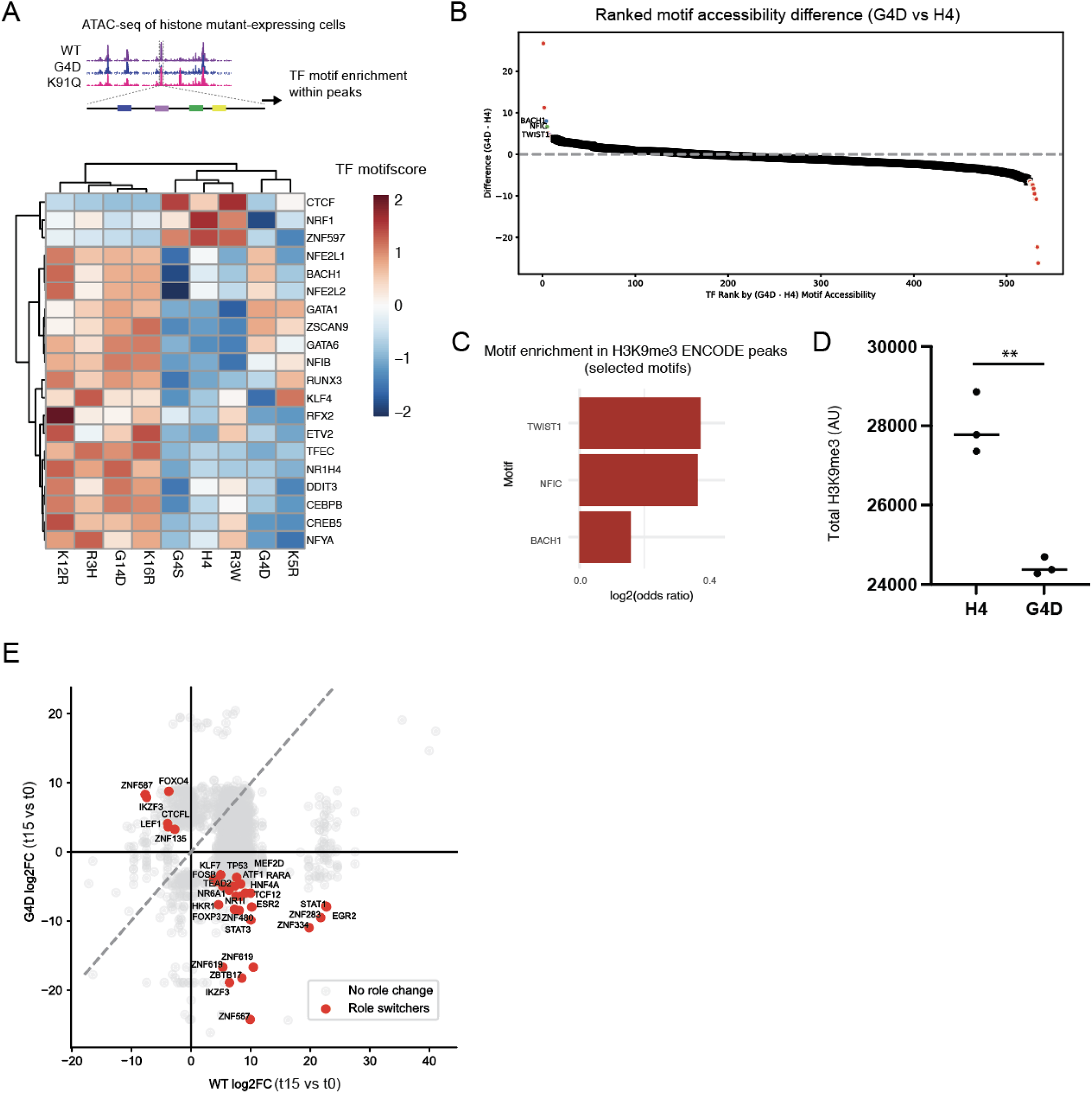
**A**. Schematic showing ATAC-seq workflow and enrichment (TF motifscore) of the most variable TF motifs in K562s. **B.** Ranked plot of motif accessibility difference in G4D vs H4 K562s. **C.** Enrichment of motifs demonstrating high accessibility increase in H3K9me3 peaks. **D.** Decrease in H3K9me3 in Jurkat T cells expressing G4D. **E.** Scatter plot of transcription factor effects on G4D vs H4 K562s, with role-switchers (significant (padj < 0.05) hits off of the diagonal) highlighted in red.

**Figure S5:**
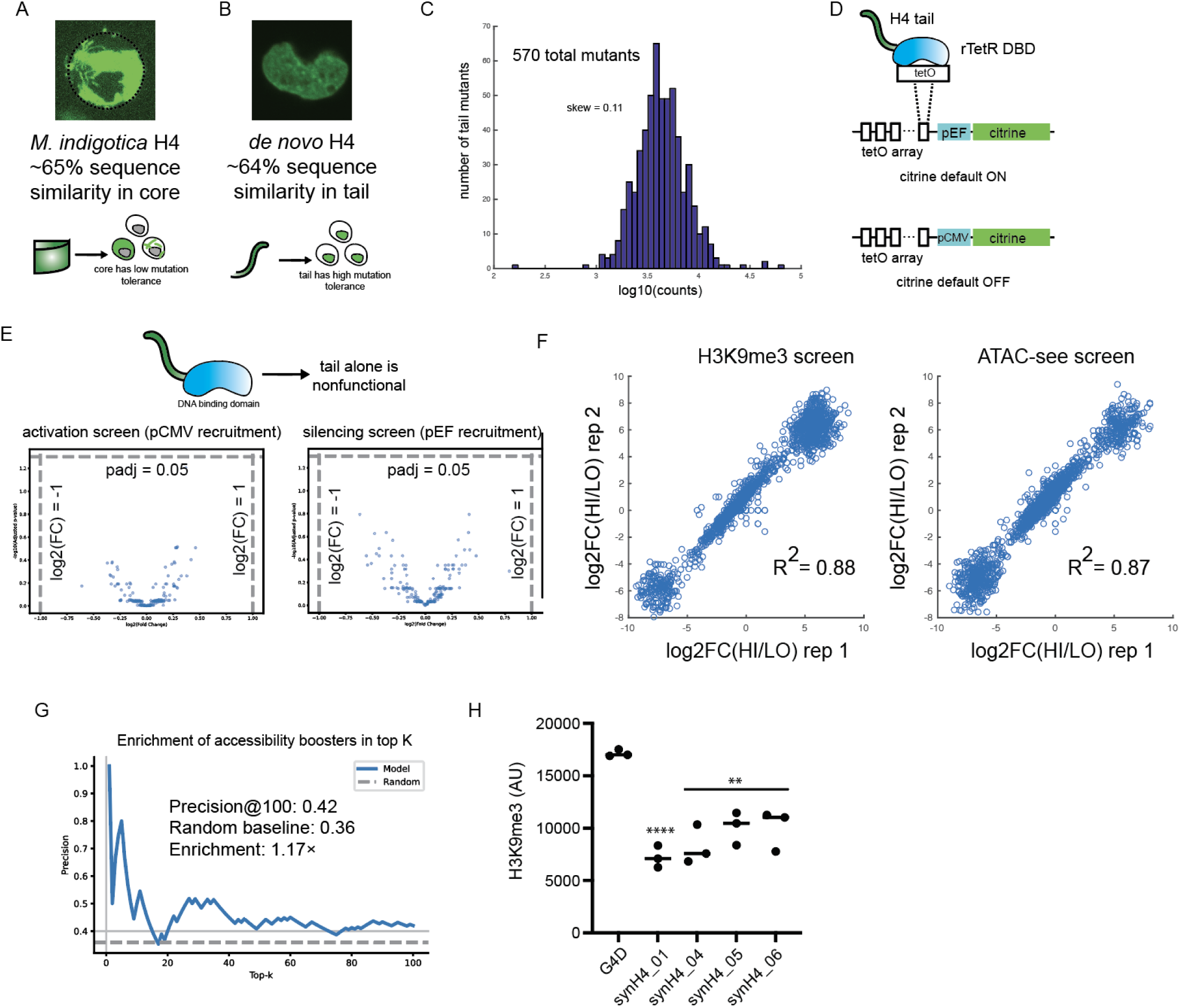
Mutations of the core (**A**) but not the tail (**B**) of H4 cause mislocalization and loss of function of the protein. **C.** Distribution of tail mutant representation over 570 total mutants, with calculated skewness, in HT-RECRUIT screens. **D.** activation and silencing HT-RECRUIT screen overview. **E.** HT-recruit screens showing lack of activity of tails alone. **F.** Replicate correlation of double-mutant H3K9me3 (left) and ATAC-see (right) screens. **G.** Precision@k plot for H4 tails that increase ATAC-see score, using the histForge model. **H.** Decrease in H3K9me3 levels in Jurkat cells expressing synH4 constructs.

